# Atlas-based classification algorithms for identification of informative brain regions in fMRI data

**DOI:** 10.1101/446856

**Authors:** Juan E. Arco, Paloma Díaz-Gutiérrez, Javier Ramírez, María Ruz

## Abstract

Multi-voxel pattern analysis (MVPA) has been successfully applied to neuroimaging data due to its larger sensitivity compared to univariate traditional techniques. Searchlight is the most widely employed approach to assign functional value to different regions of the brain. However, its performance depends on the size of the sphere, which can overestimate the region of activation when a large sphere size is employed

In the current study, we examined the validity of two different alternatives to Searchlight: an atlas-based local averaging method (ABLA, Schrouff et al., 2013a) and a Multi-Kernel Learning (MKL, Rakotomamonjy et al., 2008) approach, in a scenario where the goal is to find the informative brain regions that support certain mental operations. These methods employ weights to measure the informativeness of a brain region and highly reduce the large computational cost that Searchlight entails. We evaluated their performance in two different scenarios where the differential BOLD activation between experimental conditions was large vs. small, and employed nine different atlases to assess the influence of diverse brain parcellations.

Results show that both methods were able to localize informative regions when differences between conditions were large, demonstrating a large sensitivity and stability in the identification of regions across atlases. Moreover, the sign of the weights reported by these methods provided the directionality of univariate approaches. However, when differences were small, only ABLA localized informative regions. Thus, our results show that atlas-based methods are useful alternatives to Searchlight, but that the nature of the classification to perform should be taken into account when choosing the specific method to implement.

## 1 Introduction

The use of machine learning in neuroscience has increased exponentially in the last years, which has brought significant advances in the field (Poldrack and Farah, 2015). Hebart and Baker (2017) highlighted the existence of two independent frameworks in multivariate decoding with different aims: prediction *vs.* interpretation. Several clinical studies have employed a prediction approach, providing tools for computer-aided diagnosis of different neurological disorders, such as Alzheimer’s (Arco et al., 2015), Parkinson’s (Choi et al., 2017), epilepsy (Del Gaizo et al., 2017) or brain computer interfaces in quadriplegic patients (Blankertz et al., 2007; Nurse et al., 2015). Here, obtaining the maximum decoding performance is the main aim, whereas the source of information is not of core interest. In the interpretation context, on the other hand, machine learning is used to study the brain regions involved in different cognitive operations (Haxby et al., 2014), and here the main goal is not prediction itself. In this scenario, Multi-voxel pattern analysis (MVPA) provides larger sensitivity than classic univariate approaches (Haynes and Rees, 2006; Norman et al., 2006), as it localizes information based on the distribution of spatial patterns. Finding the most adequate analysis methods for specific contexts is of vital importance, and thus in the current investigation we compared the sensitivity of several atlas-based approaches in two different contexts to evaluate their usefulness in the field of Cognitive Neuroscience.

From the interpretation perspective, classification is simplest when performed in a region of interest (ROI) based on *a priori* knowledge. Here, the accuracy of the algorithm depends on how well the regional hypothesis fits the observed data. Haxby et al. (2001) demonstrated that the representations of faces and objects were differentially distributed in the ventral temporal cortex, whereas Haynes and Rees (2005) showed that MVPA is able to uncover the orientation-selective processing in the primary visual cortex (V1). Other studies detected distributed patterns of activity in the visual cortex (Cox and Savoy, 2003; Kamitani and Tong, 2005), whereas Poldrack (2007) highlighted the Type I error reduction when a statistical test is applied to each ROI. However, when there is not a straightforward hypothesis regarding the regions involved in specific computations, one possibility is to explore the whole brain (Mourão-Miranda et al., 2005; Balci et al., 2008; De Martino et al., 2008). The main drawback of whole-brain analyses is related to the curse of dimensionality: in fMRI studies, there are usually many more features (e.g. voxels) than samples (e.g. images or volumes). This complicates the definition of a mathematical function that separates the activation patterns related to the different experimental conditions under study (Fort and Lambert-Lacroix, 2005).

One of the most appealing approaches for the identification of informative cognitive regions is the Searchlight technique (Kriegeskorte et al., 2006), a method that offers results potentially easier to interpret due to its high spatial precision. Searchlight produces maps of accuracies from small spherical regions centered on each voxel of the brain. For each sphere, a classification analysis is performed, and the decoding performance is assigned to the central voxel. Many studies have successfully used this technique (e.g. Chen et al., 2017; Cichy et al., 2016; Coutanche et al., 2011; González-García et al., 2017; Loose et al., 2017; Qiao et al., 2017). However, it also has some disadvantages and limitations to consider. Searchlight performance depends on the size of the sphere; it usually overestimates the region of activation when a large sphere size is employed (Etzel et al., 2013; Arco et al. 2016), even when the size of informative regions stays fixed. Another drawback is that the accuracy of the classification within a certain sphere is associated with the central voxel, obviating the possibility that only a few voxels of the sphere truly contain information. Another problem is its high computational cost. Each Searchlight analysis entails a massive number of classifications, increasing the computational time compared to other simpler approaches. This time cost increases exponentially when different parameters values are evaluated (grid search) and also when permutation tests are used to evaluate the statistical significance.

There are other alternatives based on atlases that do not suffer from this large computational cost. Multiple Kernel Learning (MKL, Lanckriet et al., 2004) uses *a priori* templates of brain organization to guide the decision of the classifier. Specifically, this approach extracts information from brain parcellations provided by an atlas to maximize the performance of the classification algorithm. There are different performance measures depending on the context evaluated, from accuracy in a prediction scenario to sensitivity associated to the detection of informative brain regions in an identification context. A crucial aspect of MKL is its two-level hierarchical model. The regions used for classification have an associated weight, which indicates their contribution to the model and indexes the informativeness of the region. Voxels within each region also have a weight value. Previous studies have used this method in the context of neuroimaging, e.g. discrimination between Parkinson’s neurological disorders (Adeli et al., 2017; Filippone et al., 2013) or identification of attention deficit hyperactivity disorder (ADHD) patients (Dai et al., 2017; Qureshi et al., 2017), and localization of informative regions (Schrouff et al., 2018). MKL leads to a sparse solution, which means that only a subset of regions is selected to contribute to the decision function. However, this decreases its ability to detect informative regions. Schrouff et al. (2013a) proposed another decoding-based method based on local averages of the weights from each region defined in an atlas. This is known as Atlas-based local averaging (ABLA). First, a whole-brain classification is performed, leading to a weight map summarizing the contribution of each voxel. Then, the weights defined in each region of the atlas are averaged and normalized by the size of the region, which yields a score of the informativeness of each region. Hence, this approach builds only one classification model since the summary of the weights is done *a posteriori*. The classification model refers to the mathematical function (i.e. a hyperplane) obtained from the training procedure that separates unseen data belonging to the two different classes. These classes correspond to the activation patterns associated with the experimental conditions compared (see Section 2.3). Unlike ABLA, MKL combines the different regions of the atlas as part of the learning process: using a different atlas will result in a different classification model, with the subsequent increase in computational cost compared with ABLA.

Previous research has mostly employed atlas-based methods in prediction contexts, where the main aim is to obtain the largest accuracy possible (Arco et al., 2015; Del Gaizo et al., 2017; Illan et al., 2014). However, in interpretation contexts, employing a predefined atlas is not frequent. In this study, we evaluated the performance of different atlas-based approaches in an fMRI experiment, in two contexts with differential changes in BOLD activation. To do so, we modified the MKL and ABLA methods to better fit the requirements of an interpretation context. The standard MKL method is based on L1-regularization, a process that enforces sparsity. This means that part of the kernels (regions of the atlas) is automatically discarded for the model computation, which can be suboptimal when the main aim is to localize informative regions. For this reason, we included an L2-version of MKL (Yu et al., 2010), which avoids sparsity by allowing all regions of the corresponding atlas to contribute to the model. We compared the results obtained by MKL and ABLA methods to those obtained by Searchlight, as this approach is mainstream in current neuroimaging research. Our goal was to test alternative methods to Searchlight that overcome the limitations of this approach (dependence of the sphere size and high computational cost) while providing additional details about how information is organized in the brain. In our study, we employed nine different atlases to examine how different brain parcellations influenced the identification of informative regions of MKL and ABLA. For a contrast with large differences in the BOLD activation, we expected overlap between the significant regions obtained by all the approaches. However, this overlap would decrease for the contrast testing more subtle differences in BOLD activation. In this case, we hypothesized that the specific organization of the brain proposed by each atlas would highly affect the identification of significant regions.

## 2 Material

### 2.1 Participants

Twenty-four students from the University of Granada (M = 21.08, SD = 2.92, 12 men) took part in the experiment and received an economic remuneration (20-25 euros, depending on performance). All of them were right-handed with normal to corrected-to-normal vision, no history of neurological disorders, and signed a consent form approved by the local Ethics Committee. The sample size was chosen according to previous studies that focused on a very similar paradigm (see Gaertig et al., 2012; Chang and Sanfey, 2013 and Grecucci et al., 2013).

### 2.2 Image acquisition

fMRI data were acquired using a 3T Siemens Trio scanner at the Mind, Brain and Behavior Research Center (CIMCYC) in Granada (Spain). Functional images were obtained with a T2*-weighted echo planar imaging (EPI) sequence, with a TR of 2000 ms. Thirty-two descendent slices with a thickness of 3.5 mm (20% gap) were obtained (TE = 30 ms, flip angle = 80°, voxel size of 3.5×3.5×3.5 mm^3^). The sequence was divided in 8 runs, consisting of 166 volumes each. After the functional sessions, a structural image of each participant with a high-resolution T1-weighted sequence (TR = 1900 ms; TE = 2.52 ms; flip angle = 9°, voxel size of 1 mm^3^) was acquired.

We used SPM12 (http://www.fil.ion.ucl.ac.uk/spm/software/spm12) to preprocess and analyse the neuroimaging data. The first 3 volumes were discarded to allow for saturation of the signal. Then we used slice timing correction to account for differences in slice acquisitions. Images were realigned and unwarped to correct for head motion. Afterwards, T1 images were coregistered with the realigned functional images.

### 2.3 Design

Participants played the role of the responder in a modified Ultimatum Game (for theoretical background, which is not the focus of the present study, see Gaertig et al., 2012), deciding whether to accept or reject monetary offers made by different partners. If they accepted the offer, both parts earned their respective splits, whereas if they rejected it, neither of them earned money from that exchange. Offers consisted in splits of 10 Euros, which could be fair (5/5, 4/6) or unfair (3/7, 2/8, 1/9). The number on the left was always the amount of money given to the participant, and the one on the right was the one proposed by the partners for themselves.

Personal information about the partners was included as adjectives with different valence (16: half positive and half negative). For a third of the trials, the description were positive and another third negative, while the rest consisted on neutral trials with a text indicating absence of information (“no test”). Thus, the task contained two events in each trial, first a word (positive, negative or neutral in valence) and second two numbers (a monetary offer), to which participants had to respond. They performed a total of 192 trials, arranged in 8 runs (24 trials per run), in a counterbalanced order across participants. Each trial started with the word for 1000 ms, followed by a jittered interval lasting 5500 ms on average (4-7 s, +/0.25). Then, the numbers appeared for 500 ms followed by a second jittered interval (5500 ms on average, 4-7 s, +/0.25). Thus, participants read an adjective with a certain valence, and then they used this information to prepare to respond to the offer (second event). Thus, there is a preparatory process that leads to sustained activity along time. Because of this, the first event (words), was modelled as the duration of the word and the variable jittered interval, yielding a global duration ranging from 5 to 8 seconds. On the other hand, the second event was modelled as an impulse function (Dirac delta), i.e. with zero duration, as explained in Henson (2005), because this second event captures a different process. Once participants make a decision (accept the offer or not), the process ends. A large body of literature shows that preparatory processes extend in time (e.g. Bode and Haynes, 2009; González-García et al., 2017,2016; Sakai, 2008) whereas responding to a brief target does not (see the temporal duration of the potentials in Moser et al., 2014). This has been also measured by other neuroimaging methods, such as for example the Event-Related Potential CNV (Contingent Negative Variation, Di Russo et al., 2017).

To test the reliability of the different approaches (sensitivity and overlap of the significant regions with those obtained by Searchlight), we focused on two different classification analyses. First, we aimed at discriminating between the observed pattern associated with accepting *vs.* rejecting offers (from now on, *decision* classification). The hand used to respond was counterbalanced across participants, which means that odd subjects used the right/left hand to accept/reject an offer, whereas in even subjects the order was the opposite. Second, we focused on distinguishing the positive *vs.* negative valence of the words (e.g. Lindquist et al., 2015; from now on, *valence* classification) that were equated in number of letters, frequency of use and arousal (Gaertig et al., 2012). Extensive previous research shows that motor responses associated with accepting *vs.* rejecting offers lead to large neural differences in motor cortex (e.g. Gabay et al., 2014; Scenario 1) whereas positive *vs.* negative information leads to very subtle activation differences (e.g. Lindquist et al., 2015, Scenario 2). We employed a Least-Squares Separate (LSS) model to obtain an accurate estimation of the BOLD activation (Turner et al., 2012). This method is based on iteratively fitting a new GLM (General Linear Model, Friston et al., 1995) for each event of every trial of the experiment. The GLM is employed to compute the contribution of each experimental condition defined in the design matrix to the hemodynamic signal measured by the scanner. Thus, each model computed by LSS has two predicted BOLD time courses: one for the target event and a nuisance parameter estimate that represents the activation for the rest of the events of the same run. Previous studies have shown that this is the best approach for isolating the activation in contexts like this experiment (Abdulrahman and Henson, 2016; Arco et al., 2018), where overlap and collinearity among regressors are large.

### 2.4 Atlases

In this study, we used 9 atlases to assess the reliability of the informative regions obtained by the three atlas-based classification methods. They differ in three main aspects: the information that they use to cluster the brain regions (anatomical, functional or multimodal), the number of resulting regions (from 12 to 400) and the algorithms that implement the parcellation (a wide spectrum, from the *k*-means clustering to Bayesian models).

#### 2.4.1 BASC Cambridge

This atlas was computed from group brain parcellations generated by the BASC (Bootstrap Analysis of Stable Clusters) method, an algorithm based on *k*-means clustering to identify brain networks with coherent activity in resting-state fMRI (Bellec et al., 2010). These networks were generated from the Cambridge sample from the 1000 Functional Connectome Project (Liu et al., 2009). Based on this framework, different atlases were built depending on the number of networks defined (Urchs et al., 2015). In this study, we used four versions with 12, 20, 36 and 64 regions.

#### 2.4.2 AICHA

This atlas covers the whole cerebrum and is based on resting-state fMRI data acquired in 281 individuals (Joliot et al., 2015), and also relies on *k*-means clustering. One interesting feature is that it accounts for homotopy, relying on the assumption that a region in one hemisphere has a homologue in the other hemisphere. This leads to 192 homotopic region pairs (122 gyral, 50 sulcal and 20 gray nuclei).

#### 2.4.3 Brainnetome

Fan et al (2016) introduced an atlas based on connectivity using in vivo diffusion MRI (dMRI) and fMRI data acquired in 40 subjects. It divides the human brain into 210 cortical and 36 subcortical regions, providing detailed information based on both anatomical and functional connectivity. The number of regions was computed with a cross-validation procedure to maximize consistency across subjects (Fan et al., 2014; Liu et al., 2013). All functional data, connections and brain parcellations are freely available at http://atlas.brainnetome.org.

#### 2.4.4 Yeo2011

This atlas used a clustering algorithm to parcellate the cerebral cortex into networks of functionally coupled regions, employing fMRI data from 1000 subjects. The method employed assumes that each vertex of the cortex belongs to a single network (see Yeo et al., 2011). There are two versions available depending on the number of networks considered (7 or 17). We employed the latter for the subsequent analysis as it offers a more detailed parcellation of the brain. This atlas is preinstalled in Lead-dbs toolbox (http://www.lead-dbs.org).

#### 2.4.5 Harvard-Oxford

Clustering in this atlas was performed with the automatic algorithm presented in Desikan et al. (2006), which subdivides structural magnetic resonance data of the human cerebral cortex into gyral based regions of interest (ROI). Its validity was evaluated by computing correlation coefficients and mean distances between these results and manually identified cortical ROIs. Forty-eight cortical regions were obtained from data of 37 subjects. The resulting atlas is freely distributed with FSL (https://fsl.fmrib.ox.ac.uk/fsl/fslwiki/).

#### 2.4.6 Schaefer

This atlas adds novel parcellations and a larger precision to the brain networks published in Yeo et al. (2011) by using a local gradient approach to detect abrupt transitions in functional connectivity patterns (Schaefer et al., 2017). These transitions are likely to reflect cortical areal boundaries defined by histology or visuotopic fMRI. The resulting parcellations were generated from resting-state fMRI based on 1489 participants (see original paper for further details). There are several versions of this atlas depending on the number of regions the brain is divided into (400, 600, 800 or 1000), but we selected the first one to maintain reasonable speed on computation analyses.

## 3 Methods

In this study, we considered four different algorithms based on linear classifiers. First, the atlas-based local averaging method (ABLA) presented in Schrouff et al. (2013a). Second, an L1-MKL version of the algorithm introduced in Rakotomamonjy et al. (2008) and implemented in the PRoNTo toolbox (Schrouff et al., 2013b). Third, a modification of the L1-MKL to use an L2-norm instead of an L1 (from now on, L2-MKL) to avoid the sparsity that L1 leads to and the subsequent decrease in detecting informative regions (Yu et al., 2010). Finally, we used a Searchlight approach as a common contrast reference.

### 3.1 Atlas-based local averaging (ABLA)

This method is used after performing a whole-brain analysis in which all voxels of the brain are used as input to the classification algorithm. A linear classifier leads to a weight map where each value corresponds to the contribution of each voxel to the decision function. ABLA computes a normalization of the average weight for each region of an atlas that summarizes the importance of this region in a certain classification context. From a mathematical perspective, it is possible to specify a linear SVM (Bennett and Blue, 1998; Burges, 1998; Gaonkar et al., 2015) classification rule *f* by a pair of (***w***, ***x***), from equation:

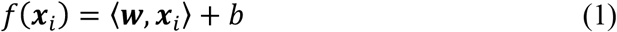

where ***w*** is the weight vector, ***x***_*i*_ is the feature vector and *b* is the error term. Thus, a point *x* is classified as positive if *f*(*x*) > 0 or negative if *f*(*x*) < 0. The decision function is based on a linear rule that maximizes the geometrical margin between the two classes, which is obtained by solving the optimisation problem described in Boser et al. (1992):

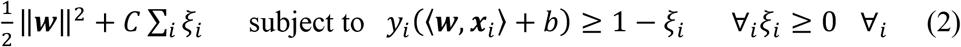

The solution to the optimization problem can be written as:

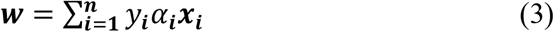

after applying the Lagrangian multipliers. Substituting the value of ***w*** in Equation 1, it is possible to rewrite the decision function in its dual form as

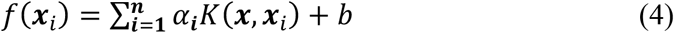

where *α*_*i*_ and *b* represent the coefficients to be learned from the examples and *K*(***x***, ***x***_*i*_) is the kernel function characterizing the similarity between samples ***x*** and ***x***_*i*_.

Once the classification model was obtained, we extracted the weight maps that guided the decision of the classifier. Then, we computed the normalized weight for each region in the atlas as the average of the absolute value of the weights contained in each region, as explained in Schrouff et al. (2013a). Equation 5 summarizes mathematically this computation:

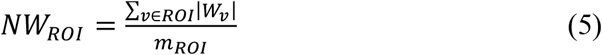

with *v* representing the index of a voxel in the weight map, *W*_*v*_ its weight and *m*_*ROI*_, the number of voxels in region ROI. Thus, the normalized weight (*NW*_*ROI*_) is a score that represents the amount of information contained in a specific brain region. A large value means that the voxels contained in the ROI have had a large contribution to the classification model.

### 3.2 Multiple Kernel Learning

Despite both ABLA and MKL rely on the brain organization provided by an atlas, they differ in the moment they use it. ABLA computes first a model from a whole-brain analysis, and then it uses the corresponding brain organization *a posteriori*. Instead, MKL combines the information from the different brain regions of an atlas to build the classification model, which means that brain parcellation is used *a priori*. Specifically, MKL combines different kernels and optimizes their contribution to the model to obtain the highest performance. As a result, this approach offers information at two levels: regions and each voxel within them. Mathematically, the decision function is computed as a linear combination of all these basis kernels as stated in Lanckriet et al. (2004):

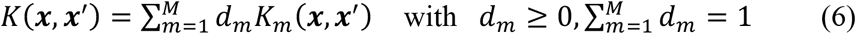

where *M* is the total number of kernels.

The decision function of the MKL problem is very similar to SVM (Equation 1) but adding the sum of the different kernels from the corresponding atlas:

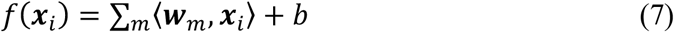

The MKL version considered in this study is based on the primal formulation of an SVM problem presented in Rakotomamonjy et al. (2008), where a solution is obtained by solving the following optimization problem:

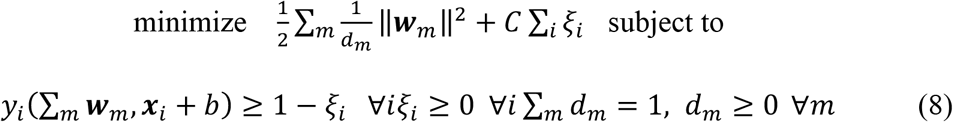

where *i* indexes the samples, *C* corresponds to the soft-margin parameter, ∑_*i*_ ξ_*i*_ is an upper-bound on the number of training errors, *b* is the bias term and *d*_*m*_ is the contribution to the decision function of each region (see Rakotomamonjy et al., 2008 for a detailed explanation).

This MKL variation optimizes, in a simultaneous manner, the contribution to the decision function of every voxel within a region and the contribution of the region as a whole, in a two-level hierarchical model. In addition, the L1-norm (Tibshirani, 1996) constraint on *d*_*m*_ enforces sparsity on some kernels. This means that several regions are automatically discarded during the learning process since a null-weight is assigned to them. Thus, from the total number of regions, only a subset of them is selected to build the classification function. This sparsity can be very interesting in prediction contexts (Arco et al., 2015; Khedher et al., 2017; Plant et al., 2010), but it can also potentiate the instability of the selected regions and decrease the sensitivity in interpretation scenarios (Baldassarre et al., 2017). For this reason, we applied a different version of MKL based on L2-norm instead of L1. In this case, sparsity is avoided, which means that all regions defined by the atlas are used to build the model. Similarly to Equation (8), the solution to the optimization problem is given by:

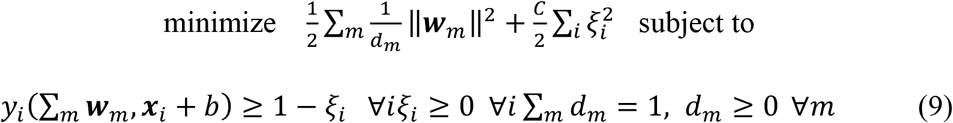

In both versions of the MKL, we applied two preprocessing steps before classification: first, we applied a mean-centering to all kernels from each region of the atlas, a very common step in machine learning. This operation relies on subtracting the voxel-wise mean for each voxel across samples, which is computed on the training data to maintain the independence between the training and test subsets. Then, we normalized the kernel dividing each sample by its norm. Regions from which kernels are computed usually have different sizes, and larger regions would have a larger contribution to the model simply because of their larger size. This operation guarantees that all regions have an equal chance regardless of their sizes.

### 3.3 Searchlight

This method was introduced by Kriegeskorte et al. (2006) to identify the location of brain regions that contain information about a given classification. It defines a sphere with a certain radius so that only the voxels inside this sphere are used to build the classification model. Performance is associated with the central voxel of the sphere. This procedure is repeated for all voxels in the brain, yielding a map of accuracies. Its main drawback is its local-multivariate nature: it extracts patterns of information from a reduced number of voxels, and this number is much smaller than the one obtained when the brain is evaluated as a whole (see Etzel et al., 2013 for additional considerations of this method).

In each sphere, we employed a support vector machine (SVM) classifier with a linear kernel due to its simplicity and the high performance reported by previous studies (Misaki et al., 2010; Pereira et al., 2009). A mathematical description of the SVM algorithm is provided in Section 3.1. We used a 12-mm radius sphere to strike a balance between sensitivity and spatial precision: smaller sizes may not detect some informative voxels whereas larger values can boost false-positives rates (Arco et al., 2016; Chen et al., 2011).

### 3.4 Performance and statistical significance

We performed a nested cross-validation to train the model and optimize the hyper-parameters of the classifier (soft-margin parameter, C), both in ABLA and in L1-MKL and L2-MKL. In these situations, the C hyperparameter range was [10^−5^:10^5^]. Regarding Searchlight, we used a standard soft-margin parameter of C=1 for each SVM classifier due to the high performance that it provides, as shown in previous studies (e.g. Chanel et al., 2016; Dosenbach et al., 2010; Fan et al., 2008). The dataset comprised an fMRI experiment divided into 8 independent runs. To maintain the independence between training and testing, we used a *leave-one-run-out* cross-validation for the external loop (all methods) and the internal loop (MKL and SVM). This means that in the Searchlight approach, 7 runs were employed to train the classifier, using the remaining one for testing. In MKL and SVM, six runs were used for training, the seventh for validation and the last one for testing. We computed the balanced accuracy within participants to evaluate the performance of the model. For a binary classification, the balance accuracy is computed as the average of the accuracy obtained in the images belonging to each experimental condition individually, which increases the robustness of the performance evaluated when there is a different number of images of each class (Brodersen et al., 2010, 2011).

Statistical significance was assessed with the method proposed by Stelzer et al. (2013), with a slight difference when the procedure was applied to Searchlight or the atlas-based approaches. Unlike Searchlight, Atlas-based methods perform a whole-brain classification, obtaining a single global accuracy. The spatial information about the amount of information in each voxel is reflected on weights. Hence, whereas in Searchlight the significance was computed from accuracy maps, in the other methods weight maps were used instead (see also e.g. Haufe et al., 2014; Schrouff et al., 2018). First, the labels of the images were randomly shuffled. Then, the corresponding classification method (ABLA, MKL or Searchlight) was applied. This procedure was repeated 100 times in a within-subject classification, resulting in 100 permuted accuracy/weight maps per participant (accuracy for Searchlight and weight for the rest). A map from each individual was randomly picked following a Monte Carlo resampling with replacement (Forman et al., 1995), averaging the permuted maps and obtaining a permuted group map. This procedure was carried out 50000 times to build an empirical chance distribution. A voxel/region was considered significant if no more than 50 samples of the empirical distribution had a larger value than the one obtained without shuffling the labels, which corresponds to a cluster-defining primary-threshold of *p*=0.001 (50/50000). Once the image was thresholded, an empirical distribution of the cluster sizes of the 50000 permuted maps was built to compute the required family-wise error rate at the cluster level. After associating a *p*-value to each cluster, a Familywise Error (FWE) correction was applied (*p*=0.05) on all-cluster *p*-values to correct for multiple comparisons at the cluster level.

### 3.5 Comparison of different atlases

Following the procedure proposed by Schrouff et al. (2013a), we computed the Pearson correlation between the overlapping voxels of the weight maps obtained by the different atlases. Since ABLA organizes the weights *a posteriori* in regions from a whole-brain classification, it is only possible to compute this correlation for L1-MKL and L2-MKL. To do so, we calculated the overlap between the significant voxels obtained by each atlas, yielding a value ranging from 0 to 1. We employed permutation tests to assess the significance of the correlation coefficients using a similar framework as described in Section 3.4.

## 4 Results

In this section, we report the results obtained by the three approaches evaluated in this study: Atlas-based local averaging (ABLA), and the two versions of Multiple Kernel Learning (L1-MKL and L2-MKL). In all cases, we only took into account results derived from above-chance accuracies that were statistically significant. Thus, we compared the weight maps of these three atlas methods that were above chance with the statistically significant accuracies map obtained by Searchlight in terms of overlap. This allows to evaluate the ability of the classification method to identify significantly informative regions regardless of the quality of the brain parcellation. Thus, the maximum overlap is determined by the overlap of the informative regions marked by Searchlight and those contained in the atlas. Moreover, for L1 and L2-MKL we show the stability of the selected regions across atlases by computing a correlation between their overlapping and statistically significant weight maps, using permutation tests to assess the significance of these correlations. We did not compute this correlation for the ABLA method because weights are exactly the same for all atlases. Additionally, we include the results obtained by these methods in two classification contexts (*decision* and *valence*) that lead to large or subtle BOLD pattern differences between the conditions contrasted, to test the generalizability of the results of the different approaches.

### 4.1 Influence of the classification methods

We first focus on comparing the results obtained by ABLA, L1-MKL and L2-MKL in the *decision* classification. Table 1 summarizes these results in terms of accuracy and overlap between the significant regions identified by each method and those obtained by Searchlight (SL). The accuracies discussed in this section correspond to the ones obtained in the maximum overlap scenario, which does not mean that these accuracies were the absolute maximum itself. We further discuss the implications of this finding in Section 5. The first approach, ABLA, yielded a maximum overlap of 70.58%, and a corresponding accuracy of 81.51%. L1-MKL led to the same maximum overlap value, 70.58%, but a higher corresponding accuracy compared to ABLA: 85.02%. On the other hand, L2-MKL obtained a maximum overlap of 77.93%, whereas the accuracy was 70.65% after employing this approach.

**Table 1:** Summary of the results obtained by the different methods and atlases in the *decision* classification. The accuracy obtained by the Searchlight approach (computed as the average accuracy of the significant voxels) is 58.79%.

| Method | Atlas | Accuracy (%) | Significant voxels | SL voxels defined in atlas | Overlap with SL (%) | Regions defined in the atlas | Significant regions |
| --- | --- | --- | --- | --- | --- | --- | --- |
| ABLA | Camb12 | 81.51 | 4704 | 4302 | 61.69 | 12 | 1 |
| L1-MKL | Camb12 | 86.2 | 4704 | 4302 | 61.69 | 12 | 1 |
| L2-MKL | Camb12 | 72.43 | 2654 | 4302 | 61.69 | 12 | 1 |
| ABLA | Camb20 | 81.51 | 3692 | 4302 | 58 | 20 | 1 |
| L1-MKL | Camb20 | 85.02 | 3692 | 4302 | 58 | 20 | 1 |
| L2-MKL | Camb20 | 65.21 | 2598 | 4302 | 58 | 20 | 1 |
| ABLA | Camb36 | 81.51 | 982 | 4302 | 21.36 | 36 | 1 |
| L1-MKL | Camb36 | 89.37 | 982 | 4302 | 21.36 | 36 | 1 |
| L2-MKL | Camb36 | 74.74 | 1000 | 4302 | 21.36 | 36 | 1 |
| ABLA | Camb64 | 81.51 | 3740 | 4302 | 63.34 | 64 | 1 |
| L1-MKL | Camb64 | 84.62 | 982 | 4302 | 21.36 | 64 | 1 |
| L2-MKL | Camb64 | 70.31 | 7613 | 4302 | 21.36 | 64 | 1 |
| ABLA | AICHA | 81.51 | 1802 | 3291 | 48 | 192 | 5 |
| L1-MKL | AICHA | 86.76 | 636 | 3291 | 19.23 | 192 | 1 |
| L2-MKL | AICHA | 69.53 | 2867 | 3291 | 60.13 | 192 | 11 |
| ABLA | Yeo2011 | 81.51 | 2731 | 3137 | 56.39 | 17 | 1 |
| L1-MKL | Yeo2011 | 87.34 | 2731 | 3137 | 56.07 | 17 | 1 |
| L2-MKL | Yeo2011 | 71.35 | 2731 | 3137 | 56.39 | 17 | 1 |
| ABLA | Harvard-Oxford | 81.51 | 4609 | 3389 | 70.58 | 48 | 2 |
| L1-MKL | Harvard-Oxford | 85.02 | 4609 | 3389 | 70.58 | 48 | 2 |
| L2-MKL | Harvard-Oxford | 70.65 | 6531 | 3389 | 77.93 | 48 | 5 |
| ABLA | Brainnetome | 81.51 | 2051 | 3057 | 47 | 246 | 9 |
| L1-MKL | Brainnetome | 78.77 | 904 | 3057 | 21.56 | 246 | 4 |
| L2-MKL | Brainnetome | 66.53 | 1129 | 3057 | 28.39 | 246 | 5 |
| ABLA | Schaefer | 81.51 | 1558 | 3137 | 42.9 | 400 | 23 |
| L1-MKL | Schaefer | 77.84 | 465 | 3137 | 14.5 | 400 | 6 |
| L2-MKL | Schaefer | 71.61 | 1926 | 3137 | 51.35 | 400 | 28 |

In the context of the *valence* classification, the ABLA method obtained a maximum overlap of 41.49%, with a corresponding accuracy of 51.77%. We assessed the significance of the accuracy by employing the non-parametric method described in Section 3.4. This last value is considerably lower than the one obtained in the *decision* classification and it likely reflects the subtle differences in the BOLD activation patterns associated with the valence of words. A complete interpretation of low classification accuracies in this kind of contexts is provided in Section 5.1. We observed that after applying the L1-MKL method, only one of the nine atlases employed led to a significant region that overlapped with Searchlight. However, the small size of this region (only 0.14% of the significant voxels obtained by L1-MKL overlapped with Searchlight results) highlights the inadequacy of L1-MKL to identify significant regions in a context like this. With reference to L2-MKL, the maximum overlap slightly increased (3.81%), with a corresponding accuracy of 49.14% (see Table 2). Since the value of the overlap is similar to the one obtained by L1-MKL, conclusions can also be applied to L2-MKL.

**Table 2:** Summary of the results obtained by the different methods and atlases in the *valence* classification. The accuracy obtained by the Searchlight approach (computed as the average accuracy of the significant voxels) is 53.74%.

| Method | Atlas | Accuracy (%) | Significant voxels | SL voxels defined in atlas | Overlap with SL (%) | Regions defined in the atlas | Significant regions |
| --- | --- | --- | --- | --- | --- | --- | --- |
| ABLA | Camb12 | 51.77 | 2095 | 911 | 41.49 | 12 | 1 |
| L1-MKL | Camb12 | 48.44 | 0 | 911 | 0 | 12 | 0 |
| L2-MKL | Camb12 | 50.71 | 0 | 911 | 0 | 12 | 0 |
| ABLA | Camb20 | 51.77 | 0 | 911 | 0 | 20 | 0 |
| L1-MKL | Camb20 | 48.18 | 0 | 911 | 0 | 20 | 0 |
| L2-MKL | Camb20 | 50.74 | 0 | 911 | 0 | 20 | 0 |
| ABLA | Camb36 | 51.77 | 0 | 911 | 0 | 36 | 0 |
| L1-MKL | Camb36 | 49.74 | 0 | 911 | 0 | 36 | 0 |
| L2-MKL | Camb36 | 49.22 | 406 | 911 | 0 | 36 | 1 |
| ABLA | Camb64 | 51.77 | 341 | 911 | 7.14 | 64 | 1 |
| L1-MKL | Camb64 | 46.88 | 549 | 911 | 0 | 64 | 1 |
| L2-MKL | Camb64 | 51.75 | 0 | 911 | 0 | 64 | 0 |
| ABLA | AICHA | 51.77 | 663 | 729 | 20.58 | 192 | 5 |
| L1-MKL | AICHA | 47.1 | 780 | 729 | 0 | 192 | 5 |
| L2-MKL | AICHA | 52.83 | 35 | 729 | 0 | 192 | 1 |
| ABLA | Yeo2011 | 51.77 | 0 | 709 | 0 | 17 | 0 |
| L1-MKL | Yeo2011 | 46.15 | 0 | 709 | 0 | 17 | 0 |
| L2-MKL | Yeo2011 | 50.97 | 1010 | 709 | 1.7 | 17 | 1 |
| ABLA | Harvard-Oxford | 51.77 | 439 | 715 | 21.4 | 48 | 1 |
| L1-MKL | Harvard-Oxford | 47.18 | 145 | 715 | 0 | 48 | 1 |
| L2-MKL | Harvard-Oxford | 48.33 | 0 | 715 | 0 | 48 | 0 |
| ABLA | Brainnetome | 51.77 | 438 | 738 | 7.99 | 246 | 3 |
| L1-MKL | Brainnetome | 43.34 | 349 | 738 | 0 | 246 | 3 |
| L2-MKL | Brainnetome | 51.19 | 137 | 738 | 0 | 246 | 1 |
| ABLA | Schaefer | 51.77 | 61 | 708 | 4.8 | 400 | 1 |
| L1-MKL | Schaefer | 46.24 | 123 | 708 | 0.14 | 400 | 2 |
| L2-MKL | Schaefer | 49.14 | 302 | 708 | 3.81 | 400 | 6 |

### 4.2 Influence of the atlases

In the first context (*decision* classification), ABLA marked as informative similar regions regardless of the atlas employed (see Figure 1). Although most of atlases led to the same results, we found a variability in terms of overlap among the different atlases (see Table 1). Specifically, the largest overlap score with Searchlight was obtained by the Harvard-Oxford atlas (70.58%), whereas the minimum value was derived from the Camb36 division of the brain (21.36%). With reference to overlap, we included the number of regions defined in the atlas and those that were marked as significant (las two columns in Table 1). For example, when ABLA was applied in combination with the Cambridge12 atlas, only 1 significant region was identified. This means that, 1/12 of the regions contained in the atlas was marked as informative in the *decision* context. Besides, 61.69% of the significant voxels obtained by ABLA overlapped with the significant voxels identified by the Searchlight, and thus ABLA missed 38.31% of the voxels identified by the Searchlight.

**Fig 1.**
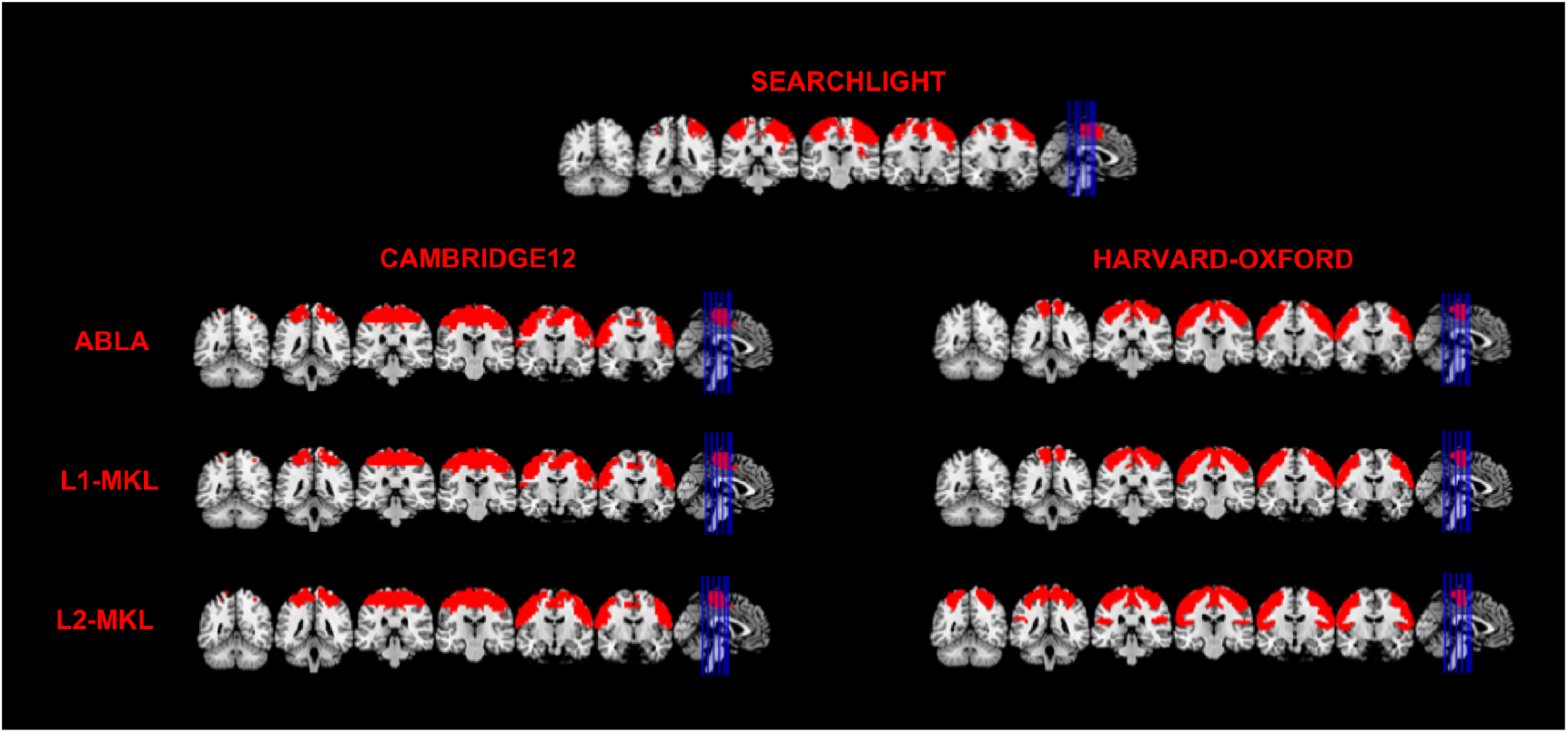
Significant voxels obtained by the Searchlight approach and the different classification methods for all the Cambridge12 and Harvard-Oxford atlases in the *decision* classification.

When applying L1-MKL in the *decision* classification, the largest overlap value was obtained by the same atlas as with ABLA: the Harvard-Oxford, with a 70.58%. However, the minimum overlap corresponded to the Schaefer atlas (14.5%). It seems that his method is more affected than ABLA by the different brain parcellations. As Figure 1 shows, the distribution of the significant regions is similar across atlases, but in this case, sensitivity is lower than ABLA for most atlases (see Table 1 for quantitative results). For the last classification method used, L2-MKL, the parcellation derived from the Camb64 atlas yielded the largest accuracy and minimum overlap score (74.74% and 21.36%, respectively). This finding highlights that maximum overlap and accuracy is not usually simultaneously obtained. The largest overlap was obtained with the Harvard-Oxford atlas (77.93%), same as when L1-MKL was used (see Figure 1 and Table 1).

Regarding the *valence* contrast, results were highly affected by the atlas used. We found a large consistency in the significant regions obtained by ABLA and Searchlight when the Cambridge12 atlas was employed. Moreover, the brain parcellations provided by AICHA and Harvard-Oxford also identified informative regions similar to those obtained by Searchlight (see Figure 2 and Table 2). Most importantly, these regions contained areas that have been reported by previous research (e.g. ventromedial prefrontal cortex, vmPFC, Lindquist et al., 2015), which supports the reliability of the results.

**Fig 2.**
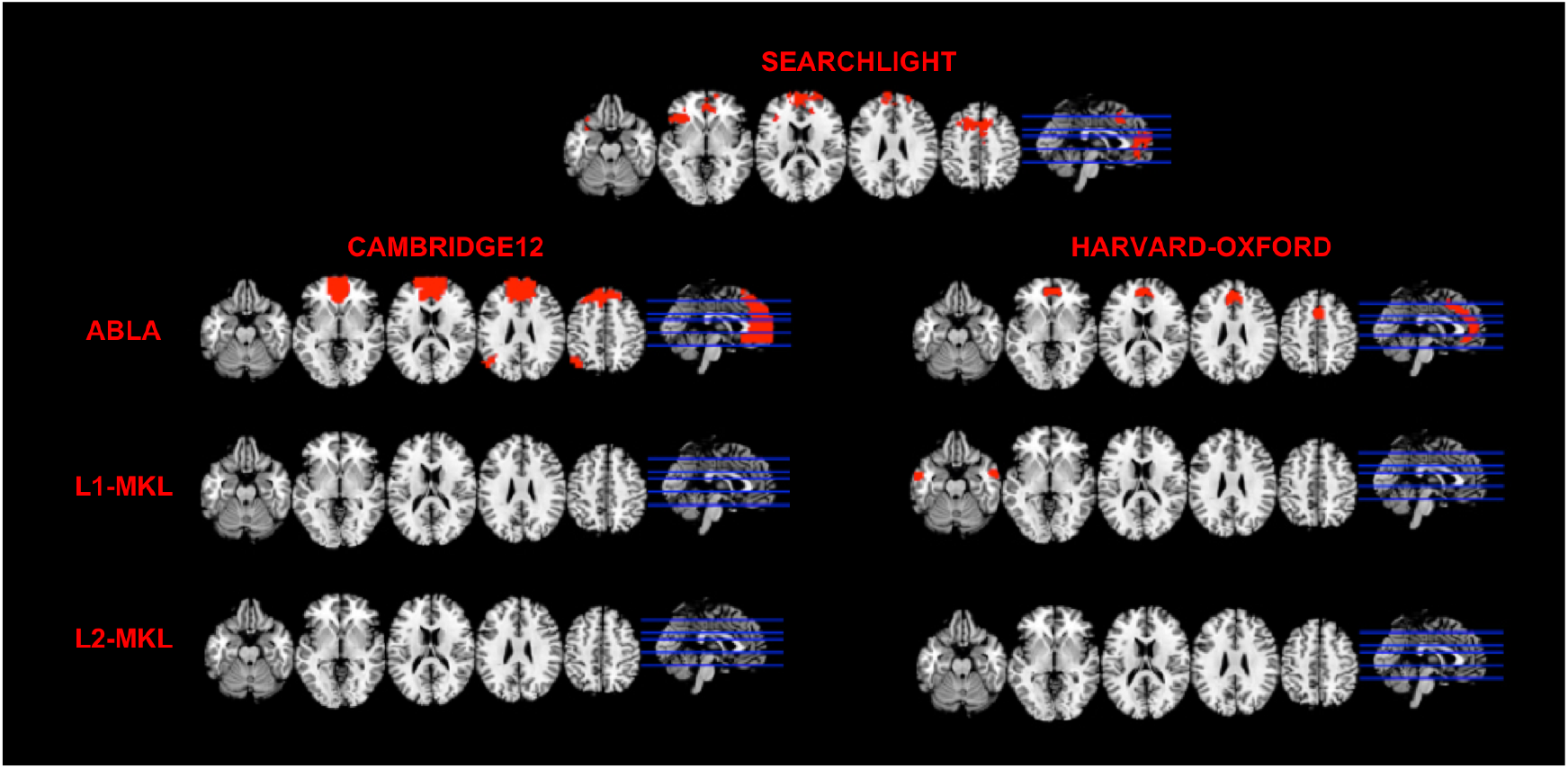
Significant voxels obtained by the Searchlight approach and the different classification methods for all the Cambridge12 and Harvard-Oxford atlases in the *valence* classification.

Unlike ABLA, the two methods based on MKL hardly detected reliable information regardless of the atlas. With reference to L1-MKL, each brain parcellation led to a completely different distribution of the informative voxels. However, none of the nine atlases that we employed yielded an accuracy above chance levels, so that the subsequent model did not provide useful information about where the information regions were located. Results were very similar for L2-MKL. Models derived from some atlases surpassed the chance level, but they were not able to identify the regions that contained information (see Figure 2 and Table 2).

### 4.3 Stability of the weights across atlases

We compared the weight maps across the different atlases for L1-MKL and L2-MKL, in the two classification contexts. In the *decision* classification, the correlation values obtained by the first 6 atlases (Camb12, Camb20, Camb36, Camb64, AICHA and Yeo2011) ranged from 0.882 to 0.975 when the L1-MKL was used. The weight maps derived from the Harvard-Oxford atlas also yielded a large similarity to these 6 atlases, but this correlation decreased when the Brainnetome atlas was employed. By contrast, the Schaefer atlas led to very different weights compared to any of the other atlases. These results suggest that, for this contrast, the decision function derived from L1-MKL is based on the same voxels. Moreover, the contribution of these voxels to the classifier decision is stable for all brain parcellations proposed by each atlas (see Table 3).

**Table 3:** Correlation between the significant weight maps across the different atlases after applying the L1-MKL method in the *decision* classification.

| L1-Multi Kernel Learning |  |  |  |  |  |  |  |  |  |
| --- | --- | --- | --- | --- | --- | --- | --- | --- | --- |
| Atlas | Camb<br>12 | Camb<br>20 | Camb<br>36 | Camb<br>64 | AICHA | Yeo2011 | Harvard-Oxford | Brainnetome | Schaefer |
| **Camb12** | 1 | 0.974 | 0.906 | 0.936 | 0.889 | 0.937 | 0.539 | 0.383 | 0.144 |
| **Camb20** | 0.974 | 1 | 0.908 | 0.951 | 0.934 | 0.947 | 0.564 | 0.552 | 0.125 |
| **Camb36** | 0.906 | 0.908 | 1 | 0.975 | 0.933 | 0.911 | 0.497 | 0.568 | 0.1 |
| **Camb64** | 0.936 | 0.951 | 0.975 | 1 | 0.963 | 0.933 | 0.542 | 0.573 | 0.081 |
| **AICHA** | 0.889 | 0.934 | 0.933 | 0.963 | 1 | 0.882 | 0.61 | 0.549 | 0.088 |
| **Yeo2011** | 0.937 | 0.947 | 0.911 | 0.933 | 0.882 | 1 | 0.566 | 0.528 | 0.193 |
| **Harvard-**<br>**Oxford** | 0.539 | 0.564 | 0.497 | 0.542 | 0.61 | 0.566 | 1 | 0.172 | 0.051 |
| **Brainnetome** | 0.383 | 0.552 | 0.568 | 0.573 | 0.549 | 0.528 | 0.172 | 1 | 0.109 |
| **Schaefer** | 0.144 | 0.125 | 0.1 | 0.081 | 0.088 | 0.193 | 0.051 | 0.109 | 1 |

L2-MKL yielded similar weight maps regardless of the atlases used (see Table 4). Only maps provided by Yeo2011 and Brainnetome are slightly less similar to those obtained by the four atlases, whereas both show a large correlation with the others. This highlights the robustness of L2-MKL in the identification of informative regions. Moreover, this finding shows the low influence that the brain parcellation has in the results, which validates the use of these atlas-based methods even without *a priori* hypothesis about the brain organization in a specific process.

**Table 4:** Correlation between the different atlases after applying the L2-MKL method in the *decision* classification.

| L2-Multi Kernel Learning |  |  |  |  |  |  |  |  |  |
| --- | --- | --- | --- | --- | --- | --- | --- | --- | --- |
| Atlas | Camb<br>12 | Camb<br>20 | Camb<br>36 | Camb<br>64 | AICHA | Yeo2011 | Harvard-Oxford | Brainnetome | Schaefer |
| Camb12 | 1 | 0.927 | 0.94 | 0.926 | 0.845 | 0.768 | 0.837 | 0.72 | 0.829 |
| Camb20 | 0.927 | 1 | 0.958 | 0.973 | 0.776 | 0.71 | 0.795 | 0.64 | 0.762 |
| Camb36 | 0.94 | 0.958 | 1 | 0.981 | 0.795 | 0.721 | 0.789 | 0.663 | 0.776 |
| Camb64 | 0.926 | 0.973 | 0.981 | 1 | 0.805 | 0.738 | 0.798 | 0.67 | 0.783 |
| AICHA | 0.845 | 0.776 | 0.795 | 0.805 | 1 | 0.948 | 0.94 | 0.904 | 0.973 |
| Yeo2011 | 0.768 | 0.71 | 0.721 | 0.738 | 0.948 | 1 | 0.897 | 0.857 | 0.945 |
| Harvard-Oxford | 0.837 | 0.795 | 0.789 | 0.798 | 0.94 | 0.897 | 1 | 0.849 | 0.945 |
| Brainnetome | 0.72 | 0.64 | 0.663 | 0.67 | 0.904 | 0.857 | 0.849 | 1 | 0.889 |
| Schaefer | 0.829 | 0.762 | 0.776 | 0.783 | 0.973 | 0.945 | 0.945 | 0.889 | 1 |

For the *valence* classification, in contrast, the localization of the informative regions was so variable that results derived from most atlases did not overlap. For this reason, we could only compute the correlation between AICHA, Harvard-Oxford and Brainnetome for L1-MKL, which yielded a maximum overlap of 0.428 (see Table 5). Results obtained by L2-MKL also showed a reduced overlap between the weight maps and we could only correlate the significant results of AICHA, Yeo2011 and Schaefer. In this case, the maximum correlation was obtained by Yeo2011 and Schaefer, yielding a value of 0.99 (see Table 6). Nevertheless, this value was obtained from a small region since the spatial distribution of the significant results provided by these two atlases were considerably different.

**Table 5:** Correlation between the significant weight maps across the different atlases after applying the L1-MKL method in the *valence* classification.

| L1-MKL |  |  |  |  |  |  |  |  |  |
| --- | --- | --- | --- | --- | --- | --- | --- | --- | --- |
| Atlas | Camb<br>12 | Camb<br>20 | Camb<br>36 | Camb<br>64 | AICHA | Yeo2011 | Harvard-Oxford | Brainnetome | Schaefer |
| **Camb12** | 1 | - | - | - | - | - | - | - | - |
| **Camb20** | - | 1 | - | - | - | - | - | - | - |
| **Camb36** | - | - | 1 | - | - | - | - | - | - |
| **Camb64** | - | - | - | 1 | - | - | - | - | - |
| **AICHA** | - | - | - | - | 1 | - | 0.217 | 0.428 | - |
| **Yeo2011** | - | - | - | - | - | 1 | - | - | - |
| **Harvard-**<br>**Oxford** | - | - | - | - | 0.217 | - | 1 | - | - |
| **Brainnetome** | - | - | - | - | 0.428 | - | - | 1 | - |
| **Schaefer** | - | - | - | - | - | - | - | - | 1 |

**Table 6:** Correlation between the significant weight maps across the different atlases after applying the L2-MKL method in the *valence* classification.

| L2-MKL |  |  |  |  |  |  |  |  |  |
| --- | --- | --- | --- | --- | --- | --- | --- | --- | --- |
| Atlas | Camb<br>12 | Camb<br>20 | Camb<br>36 | Camb<br>64 | AICHA | Yeo2011 | Harva<br>rd-Oxford | Brainnetome | Schaefer |
| **Camb12** | 1 | - | - | - | - | - | - | - | - |
| **Camb20** | - | 1 | - | - | - | - | - | - | - |
| **Camb36** | - | - | 1 | - | - | - | - | - | - |
| **Camb64** | - | - | - | 1 | - | - | - | - | - |
| **AICHA** | - | - | - | - | 1 | - | - | - | 0.975 |
| **Yeo2011** | - | - | - | - | - | 1 | - | - | 0.99 |
| **Harvard-**<br>**Oxford** | - | - | - | - | - | - | 1 | - | - |
| **Brainnetome** | - | - | - | - | - | - | - | 1 | - |
| **Schaefer** | - | - | - | - | 0.975 | 0.99 | - | - | 1 |

### 4.4 Directionality of the weights

In the *decision* classification, we evaluated not only the localization of the informative weights but their sign. Due to the nature of the contrast (odd participants used their right hand to accept an offer, whereas even participants employed their left hand), it was expected that weights were organized according to their sign in a specific hemisphere for each group of participants. Figure 3 shows the distribution of the significant voxels depending on the sign of their weights for the ABLA method, comparing them with results obtained by univariate analysis. It is remarkable that participants who accepted the offer with the right hand and rejected it with the left hand (odd group) show a cluster of positive weights in the left hemisphere and a cluster of negative weights in the right hemisphere. On the other hand, these results are shifted when results from even participants were evaluated: the right hemisphere contains weights associated with accepting an offer, whereas the left hemisphere shows the negative weights. These results are consistent with those obtained by univariate analysis. Results for both MKL methods are very similar to the ones obtained by ABLA. For those participants that accepted the offer by employing their right hand, the weights in the right hemisphere are positive, whereas the same hemisphere in the group of people that used their left hand to accept the offer contains negative weights (see Figure 3).

**Fig 3.**
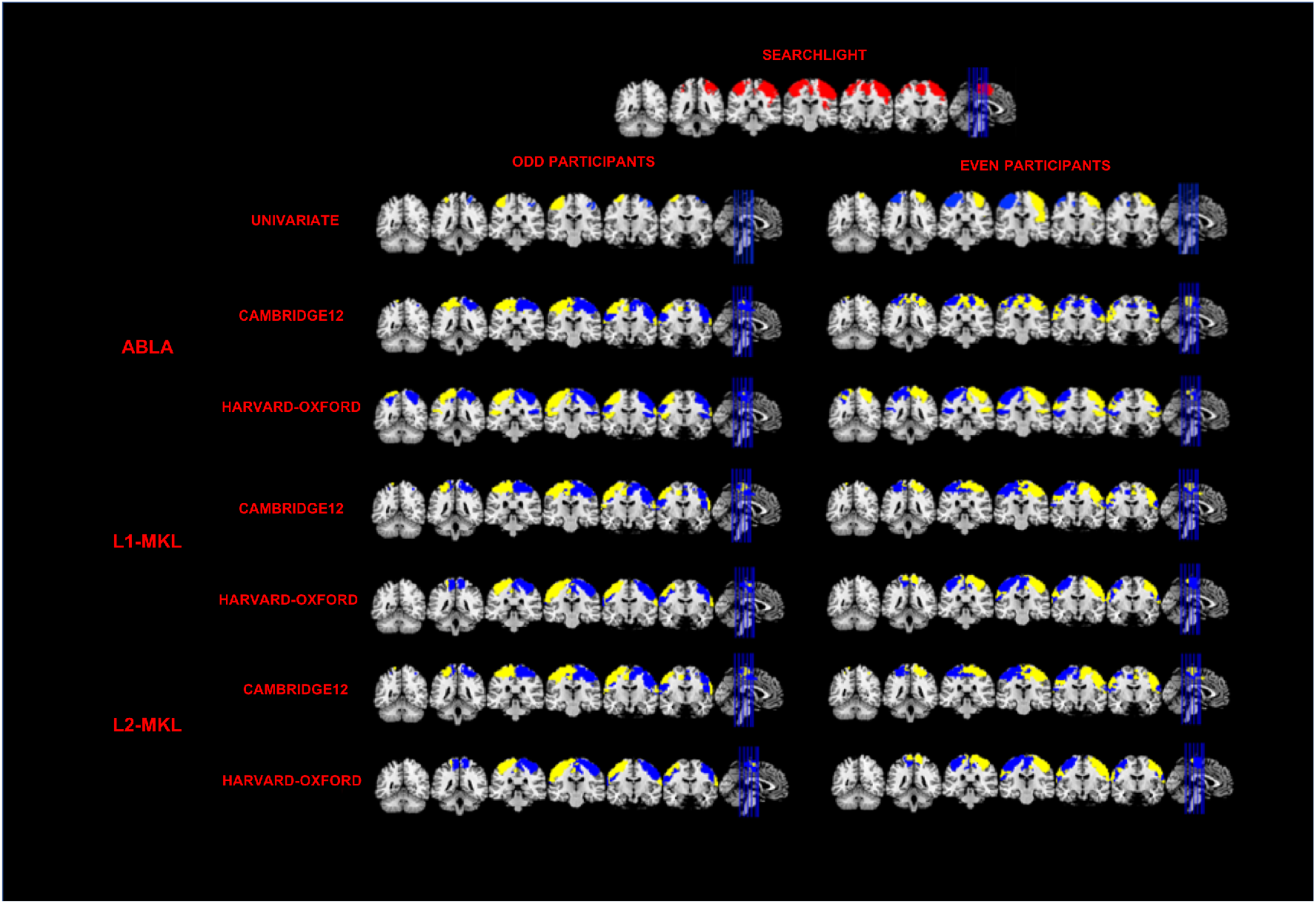
Summary of the results obtained for the *decision* classification by Searchlight, ABLA, L1-MKL, L2-MKL and univariate approaches. The three classification methods show large differences between the two groups considered (odd/even participants). Searchlight only provides information about the significance of each voxel itself, so that no separation between groups was considered.

## 5 Discussion

In this study, we evaluated atlas-based methods, alternative to Searchlight, to localize the informative regions involved in cognitive functions. We extracted the weight maps from three atlas-based classification approaches (ABLA, L1-MKL and L2-MKL) and evaluated the statistical significance of each region. We used these methods in two different contexts. In the first one, where the two classes generated large differences in the observed pattern, L2-regularization resulted the best option for interpretation purposes. Moreover, atlas-based approaches showed a large stability in the informative regions found regardless of the atlas employed, which highlights the adequacy of these methods. In contrast, when the differences in the BOLD patterns associated with each class were subtle, only the ABLA approach showed certain stability in the informative regions across the atlases. In what follows we discuss the implications for choice of classification methods, atlases, and the role of the weights.

### 5.1 Influence of the classification methods

Our results indicate that maximum accuracy and overlap do not usually concur, especially when only subtle differences exist between the patterns. In the *decision* classification, we found differences across the methods in terms of overlap and accuracy. We can separate the different approaches in two groups: on the one hand, ABLA and L2-MKL; on the other, L1-MKL. They differ in the way regularization is performed: while ABLA and L2-MKL use an L2-norm regularization, L1-MKL employs an L1. The dimensionality reduction provided by the L1-norm can be helpful from the prediction standpoint given the larger accuracies obtained. However, our results show that the model with the largest overlap is not usually the most accurate, which is consistent with previous studies (e.g. Sona et al., 2007). Our results stress the need of clearly separating the use of multivariate decoding for prediction and for interpretation (Hebart and Baker, 2017), and highlight the importance of selecting the methods that best fit the desired aim.

In the *valence* classification, we found larger differences across the methods in terms of overlap and accuracy. ABLA was the approach that obtained the largest overlap with Searchlight results, whereas L1-MKL and L2-MKL hardly detected significant regions. The key of this finding is the classification problem itself. Isolating regions with a differential involvement in valence processing is a difficult endeavor, as shown by recent metanalytic approaches (Lindquist et al., 2015). Moreover, previous research has shown that decoding accuracies in the prefrontal cortex (PFC) are much lower than in other regions like the visual cortex (e.g. see Figure 1 in Bhandari et al., 2018), similar to the accuracy obtained by ABLA in the *valence* classification.

The values of the classifier’s accuracy are influenced by how information is represented in the brain and the sensitivity of the neuroimaging technique employed. As an example, single-unit recordings have demonstrated that although face identity is represented in underlying neuronal populations as measured by electrophysiological single-unit recordings, this information cannot be retrieved from fMRI data, due to the lower spatial resolution of the neuroimaging method (Dubois et al., 2015). Accuracies can be theoretically relevant once they are statistically significant above-chance levels, regardless of their value (Hebart and Baker., 2017; Bhandari et al., 2018). We employed a very stringent significance threshold (*p*<0.001) and an FWE correction was applied to the resulting p-values to address the multiple comparisons problem. Thus, although some accuracies have low values, only those that are statistically significant (i.e. consistent above-chance across participants) are included as informative.

Our results show that ABLA provides a larger overlap than the MKL methods in the two classification problems, especially in the *valence* one. This discrepancy must be due to the different framework of ABLA. MKL approaches use the regions of the atlas to build the model during the learning process. This means that the atlas should properly delimitate the different regions involved in the context under study. Otherwise, the resulting model would be suboptimal. Instead, ABLA builds the classification model from a whole-brain parcellation, incorporating the brain organization afterwards. For this reason, ABLA leads to a better performance in conditions of subtle or small differences between the experimental conditions. However, MKL methods would be more sensitive when the atlas leads to a realistic approximation of the brain subdivisions.

It is worth mentioning that regions that Searchlight marked as informative in the two classification contexts are conceptually logic and replicate previous results in the literature (Gabay et al., 2014; Kuzmanovic et al., 2018; Lindquist et al., 2015). This means that Searchlight results are accurate enough to be used as a reference of how informative regions are spatially distributed. Thus, computing the overlap between the atlas-based classification approaches and Searchlight is an optimal way of evaluating the ability of the first methods to identify informative brain regions.

### 5.2 Influence of the atlases

Results show that the specific brain parcellation of each atlas impacts the spatial accuracy of the different methods only when differences in the observed pattern are small. In the *decision* classification, our results evidence that informative regions can be identified even when the brain parcellations provided by the atlases do not perfectly delimitate the regions involved in the cognitive function under study. However, in contexts like the *valence* classification, these atlases are not accurate enough to guarantee the identification of the sources of information. This is probably related to the size and the specific shape of the region involved in a certain cognitive function, such as the vmPFC associated with the *valence* classification. The only region that ABLA marked as significant in the Camb12 parcellation is the one that contains the vmPFC, which has a massive size in this atlas. Since atlas-based methods consider each region as a whole, a large number of voxels are marked as significant only because they are in the same region as the ones that are really informative. Here, the parcellation proposed by the atlas is a good match to the spatial organization of a structure such as the vmPFC, leading to a higher sensitivity and spatial accuracy.

The number of subdivisions of the atlases also influenced the performance of the algorithms employed. In the *decision* classification, the optimum value in terms of overlap was obtained by the 36 regions that the Camb36 atlas is divided into. Using an atlas with few subdivisions means that it is more likely to find an informative region. Instead, a large number of parcellations means that the classifier has to be much subtler in the identification of informative regions. The parcellations derived from Schaeffer add larger precision and subdivisions to the brain networks published by the Yeo2011 atlas. However, results show a better performance in terms of sensitivity when the simplest one was used. This strongly indicates that using atlases that contain large regions is similar to employing large Searchlight spheres where only a few voxels within are informative (Etzel et al., 2013). This increases the probability of marking as significant voxels that are not, increasing the false-positives rate.

### 5.3 Stability of the weights across atlases

We have found a large correlation between the significant weight maps obtained by different atlases in the *decision* classification. For the L1-MKL approach, all atlases except Schaefer led to large correlation values. Hence, the weights associated with each model were very similar, which highlights the stability of the classification methods. Interestingly, we found the largest correlations in the weight maps obtained by the four Cambridge atlases, which are derived from the same clustering algorithm (BASC). This supports the idea that the mathematical framework employed to delimitate the different brain regions in an atlas can influence the success of the subsequent analyses. On the other hand, the poor performance of L1-MKL when the Schaefer atlas was used can be due to the conjunction of a sparse method and an atlas with a large number of regions. It is important to note that our results do not invalidate the use of ambitious atlases aiming at obtaining a detailed parcellation of each cortical region. However, if these parcellations do not match how information is represented in the brain, sparse solutions are not recommended. Unlike L1-MKL, L2-MKL obtained a large correlation score between each pair of atlases (Table 4): the weight maps that guide the classification are essentially the same regardless of the atlas used. This evidences that it is possible to employ this approach even without a clear hypothesis about the brain organization in a specific context.

Nevertheless, these conclusions are only valid when there are large differences in the observed pattern associated with the two classes to distinguish from. Our findings in the *valence* classification differ substantially from those obtained in the *decision* classification. L1-MKL results (summarized in Table 5) show that we could hardly compute the correlation between two pairs of atlases: the first one, AICHA and Harvard-Oxford; the second, AICHA and BN. In addition, none of the significant results provided by these atlases share any voxel with the Searchlight results, which illustrates that weight maps are similar from a mathematical perspective, but make a null contribution to the neuroscience standpoint. Results obtained by L2-MKL are summarized in Table 6 and conclusions derived from them are essentially the same than L1-MKL. We could only compute the correlation between two pairs of atlases: Schaefer-AICHA and Schaefer-Yeo2011. From these three atlases, Schaefer is the one that leads to some overlap with the Searchlight approach: 3.81%. However, none of these significant voxels are shared by AICHA and Yeo2011. This reflects that the two versions of MKL are not able to identify small informative regions in contexts where differences in the observed patterns are minimum. According to these results, investigators should use ABLA when their paradigm produces a small difference in BOLD activation patterns. However, the performance of the MKL approaches could increase if the brain parcellations derived from an atlas adapts to the concrete pattern of activations obtained in a particular task. This could be done by employing methods based on machine learning to cluster the different brain regions from individual fMRI data of each subject (Gordon et al., 2017; Schaefer et al., 2018; Wang et al., 2015), which could boost the spatial precision in the brain parcellations. Thus, it is likely that L2-MKL could obtain similar or even better results than ABLA in this scenario, but future studies should assess this.

### 5.4 Directionality of the weights

One of the main advantages of using weights instead of accuracy is the directionality that they provide. The term directionality is related to univariate analysis, which localize the regions where the activation associated with an experimental condition is larger than the activation associated with another condition. We have evaluated the sign of the weight of each voxel within the significant regions for the three atlas-based methods for the *decision* classification. The three approaches obtained a map in which weights were organized according to their sign. For odd participants, regions associated with the acceptance of an offer (use of the right hand) were localized in the left hemisphere, with a positive sign. On the other hand, regions that contained information when the offer was rejected (left hand) were found in the right hemisphere, with a negative weight. More importantly, the informative regions for even participants shifted: positive weights were found in the right hemisphere, whereas weights with a negative sign were found in the left hemisphere.

The large similarity between these results and those obtained by the univariate approach (see Figure 3) is remarkable. Regions with a larger activation when participants accept/reject an offer match the sign of the weights of the different multivariate methods. However, atlas-based approaches use normalized data, which eliminates the differences in the global activation levels associated to each condition. Thus, these methods identify areas that show a different spatial distribution of the information, while the univariate approach purely relies on differences in the activation level.

Another difference between multivariate and univariate methods is the different sensitivity that they offer. We employed a classification context where differences at the cluster level could also be picked up by univariate methods to highlight one of the main advantages of atlas-based approaches. Similarly to Searchlight, these techniques are able to extract information from fine-grained differential activation patterns, which results in a boost in sensitivity compared to univariate analysis. Figure 3 reveals the differences in sensitivity between multivariate and univariate methods in the *decision* context, whereas differences between these two approaches in the *valence* classification can be found in previous studies (see Figure 10 in Arco et al., 2018 and Figure 1 in Lindquist et al., 2015). Thus, employing weights provides the sensitivity of multivariate approaches and the directionality of univariate ones. This is quite useful from a Cognitive Neuroscience point of view, as it allows not only to detect the brain regions that contain information about a specific cognitive process, but also to identify how this information is distributed.

## 6 Conclusions

In this study, we compared three different atlas-based approaches to Searchlight to assess their ability to identify informative brain regions for cognitive contrasts that generate either large or small differences in BOLD activation patterns. We have shown for the first time that these methods can be used as an alternative to Searchlight since they localize informative regions when there are large differences between the observed patterns associated with the two classes to distinguish from. In this case, results are consistent across atlases, which manifests that these approaches can be used even without a prior hypothesis about the concrete pattern of activations expected. Moreover, the use of weight maps provides additional relevant information by combining the sensitivity of decoding analyses and the directionality of univariate approaches. This is extremely interesting in interpretation scenarios, where the main goal is to localize informative regions and to identify how information is distributed. However, results in terms of sensitivity change drastically when the differential observed pattern is much lower. Methods based on MKL are highly affected by the discrepancy of the shape of brain regions containing information and the one proposed by the atlases. On the other hand, ABLA is the only approach that identifies informative regions in accordance with previous research, which means that it is the most trusted method when subtle differences are evaluated. Our results pave the way for finding a method that leads to a large spatial accuracy in the identification of subtle changes of the observed patterns. Future studies are needed to evaluate the performance of these methods when the brain parcellations are specifically computed for each participant, which may substantially improve the neuroanatomical functional precision and the subsequent identification of informative regions.

## 7 Funding

This work was supported by the Spanish Ministry of Science and Innovation through grant PSI2016-78236-P to M.R and the Spanish Ministry of Economy and Competitiveness through grant BES-2014-069609 to J.E.A.

## 8 Acknowledgments

We are grateful to Janaina Mourão-Miranda for her kind help during the development of the algorithms employed in the current research.

## 9 Conflict of Interest

The authors declare that the research was conducted in the absence of any commercial or financial relationships that could be construed as a potential conflict of interest.

